# Divergent Behavior Amid Convergent Evolution: A Case of Four Desert Rodents Learning to Respond to Known and Novel Vipers

**DOI:** 10.1101/362202

**Authors:** Sonny S. Bleicher, Burt P. Kotler, Omri Shalev, Dixon Austin, Keren Embar, Joel S. Brown

## Abstract

Desert communities word-wide are used as natural laboratories for the study of convergent evolution, yet inferences drawn from such studies are necessarily indirect. Here, we brought desert organisms together (rodents and vipers) from two deserts (Mojave and Negev). Both predators and prey in the Mojave have adaptations that give them competitive advantage compared to their middle-eastern counterparts. Heteromyid rodents, kangaroo rats and pocket mice, have fur-lined cheek pouches that allow the rodents to carry larger loads under predation risk compared to gerbilline rodents. Sidewinder rattlesnakes have heat-sensing pits, allowing them to hunt better on moonless nights when their Negev sidewinding counterpart, the Saharan horned vipers, are visually impaired. In behavioral-assays, we used giving-up density (GUD) to gage how each species of rodent perceived risk posed by known and novel snakes. We repeated this for the same set of rodents at first encounter and again two months later following intensive “natural” exposure to both snake species. Pre-exposure, all rodents identified their evolutionarily familiar snake as a greater risk than the novel one. However, post-exposure all identified the heat-sensing sidewinder rattlesnake as a greater risk. The heteromyids were more likely to avoid encounters with, and discern the behavioral difference among, snakes than their gerbilline counterparts.

## Introduction

Deserts, and desert rodents in particular, provide a model system for studying parallel and convergent evolution. Deserts around the world form at least five evolutionarily independent laboratories of adaptation, ecology, and evolution [1–6]. Shared environmental conditions of temperature, precipitation, and aridity force evolutionary processes in a manner that results in similar adaptations in species that fill similar ecological roles. Not only do species converge, but communities may too [7–10]. A good example of this can be studied in desert dunes of the Mojave and of the Negev deserts. In both of these systems we find an array of plants that drop their seeds onto the sand (creating a seed bank); a variety of rodent species feed on these seeds [11–13]; and medium-sized sidewinding vipers feed on the rodents [14,15].

The Mojave and Negev deserts of North America and the Middle East, respectively, possess rodents with similar ecologies [5,7,13,16,17]. These rodents are nocturnal, semi-fossorial, seed-eating, and seed caching. However, the heteromyid rodents of the Mojave may have a constraint breaking adaptation compared to their convergent counterparts in the Negev, the gerbilline rodents. A constraint-breaking adaptation is a game-changing evolutionary adaptation that alters, relaxes or eliminates tradeoffs and confers a competitive advantage to its holder over those lacking the trait as defined by Rosenzweig and McCord [18]. The heteromyids have external fur lined cheek pouches that allow them to stow large quantities of food (in good dry conditions) before having to return to a burrow for caching [11]. In contrast, the gerbilline rodents carry their grain in their mouths, logically requiring more trips to collect the same quantity of grain and thus increasing exposure to predators.

Similar to the rodents, rattlesnakes from North America and horned vipers from the Middle East provide a textbook example of convergence [19]. Despite being separated by 18 million years from their most recent common ancestor [20], each has evolved the same locomotion method, similar coloration patterns, and a similar adaptation of scales over the eye ridge, protruding as horns. However, the North American sidewinder belongs to the evolutionary lineage of pit-vipers, a lineage that evolved infra-red heat sensing pits. The pit-vipers provide another example of a constraint breaking adaptation compared with Saharan horned vipers. The heat-sensing pits enable the sidewinder to be active on dark nights with no ambient moonlight. The pits also enable safer, more precise strikes at warmer, more vulnerable locations in their endothermic prey [21].

We report here an intercontinental comparison for how two species of Mojave Desert rodents and two species of Negev Desert rodents respond to their evolutionarily and ecologically familiar versus novel snakes. We ask three questions: (1) Do gerbils and heteromyids assess risk from snakes in a similar manner? That is, do they make the same choices when facing snakes with and without heat-sensing pits? (2) Do both sets of rodents assess risk from novel predators as equal to that of evolutionarily familiar ones? (3) Does a prolonged (two-month) exposure to both snakes (in a larger and more realistic setting) diminish the perceived risk of predation from horned vipers, compared with heat-sensing pit-vipers? If so, do both sets of rodents reach the same conclusion, i.e. exhibit the same behavioral response?

## METHODS

### Study Species

We brought together one large and one small coexisting desert rodent from each continent, two common gerbils from the Negev Desert of Israel and a kangaroo rat and a pocket mouse from the deserts of the southwestern United States, to a common and controlled setting in the Negev Desert. The Negev Desert gerbils include the greater Egyptian gerbil [GP] (*Gerbillus pyramidum*), 40 g, and Allenby’s gerbil [GA] (*Gerbillus andersoni allenbyi*), 30 g [22]. The North American Desert rodents include Merriam’s kangaroo rat [DM] (*Dipodomys merriami*), 45 g [23], and the desert pocket mouse [CP] (*Chaetodipus penicillatus*), 22 g [24]. All are nocturnal desert granivores commonly found on sandy substrates such as sand dunes. All four rodents have adaptations to reduce the risk of predation, including saltatorial locomotion for enhanced escape abilities and auditory adaptations to increase hearing acuity. These adaptations are especially well developed in the kangaroo rats [4,25].

We brought wild-caught vipers, trapped at locations where they would come in contact with wild populations of the above-mentioned rodents, to the same facility. We caught sidewinder rattlesnakes (*Crotalus cerastes)*, 35-60cm mean length, from the Mojave Desert [26] and Saharan horned vipers (*Cerastes cerastes*), 30-60cm mean length, from the Negev Desert [15]. Both snakes side-wind, borrow in the sand (usually under bushes) and feed on a variety of rodents and lizards [14,15].

Animal collection was done respectively in the Mojave and Negev Deserts. The heteromyids were predominantly trapped in the Parker Dunes area (N 34°9’7.969”, W 114°7’34.245”) and supplemented by individuals from the San Bernardino (AZ) area (N 31°23’22.082”, W 109°11’ 22.851”). The sidewinders were collected in the Avra Valley alongside country roads (N 32°24’49.335”, W 111°29’38.138”). The gerbils in Israel were collected in the Mashabim Dunes (N 31°0’14.531”, E 34°44’47.31”) and the horned vipers on the border between Israel and Egypt at Be’er Milka (N 30°57’4.609”, E 34°23’10.821”).

### Experimental Design

We used an “interview” approach [27–29] to measure the response of each rodent species to the risks posed by the two snake species. We measured the response prior to exposure of the rodents to the novel viper species and following a two-month exposure to both snake species in an semi-natural arena (described in Bleicher 2014; Bleicher et al. 2016). We assessed the response of each species to the vipers using a metric borrowed from foraging theory, the giving-up density (GUD; Brown 1988). The GUD is the amount of food a forager leaves behind untouched in a resource patch to measure foraging efficiencies and costs. Most relevant for our purposes is that these costs include those arising from the perceived risk of predation. Hence the forager will leave a lower GUD when it perceives lower risk [31].

The experiments were conducted in a light-controlled room at Ben Gurion University of the Negev’s Blaustein Institutes for Desert Research, at Midreshed Ben Gurion, Israel (N 30°51’17.401”, E 34°47’6.637”). We erected a total of eight, 3-compartment (henceforth room) behavioral-assay systems (henceforth system), which we call interview chambers (*S1*). We called them interview chambers as they allow the researchers to question an individual animal and allow the animal to rate how it perceives treatments in relation to each other. With repeated measures it allows the researchers to obtain how each individual changed its perception of the controlled treatments, *i.e*. it’s opinion, after a manipulation.

Each system consisted of a circular nest-box attached to three 80 x 40 x 40 cm test-rooms. Each room was connected to the nest-box with a 30 cm PVC tube to allow rodents free movement between the nest-box to any room. Each room was large enough to contain a small cage and a 38 x 28 x 8 cm foraging tray (Sup. Appendix 1). Each foraging tray was set with 2 liters of sand and 1.5g of millet. For each system during a trial, the cage in one compartment contained a sidewinder rattlesnake, the cage in the second contained a horned viper, and the cage in the third was empty. In order to avoid the possibility of directional bias, we randomized the positions of treatment-rooms (henceforth treatment) to different cardinal directions in each system.

between dusk and dawn we conducted a maximum of five, 2-hour tests (henceforth round), in which a single rodent was placed in the nest-box and was allowed to forage in the system. This allowed sufficient time for rodents to move among compartments and forage in trays under the different treatments. Each rodent was run at random times each night to nullify the preferred activity periods of the rodents. The rodents weren’t fed prior to the experiment adding incentive to forage when in the chambers. Following the experiment, animals were returned to their holding cages and fed 3 grams of millet and a mealworm (*Tenebrio molitor*) to offset stress related calorie loss. Following each test, each of the foraging trays was sieved and the remaining millet removed and weighed to obtain the GUD. The systems were reset after each rodent was tested with fresh millet and the next rodent introduced for the following round.

Each individual rodent was tested for two nights pre-exposure and an additional two nights post-exposure, with a night between runs to avoid possibly over-stressing the animals. The exposure periods are two-month experiments in which the rodents cohabitated in a semi natural arena (aviary dimensions 17 x 34 m) with uncaged snakes of both species (3 of each species) allowing them to learn the differences in behavior of the predators. In addition, by flying an owl in the aviary on half the nights, we compared how the risk from snakes compares to the risk from owls (*cf*. Bleicher 2014). Our aviary provides a system where the rodent enjoy ideal free distribution through special gates within the arena, but predators are limited in movement by the same gates. Using RFID tags implanted subcutaneously in the rodents, and loggers under food patches we are certain the majority of rodents experienced encounters with both snake species. We pre-{post-} interviewed 51 {19} GAs, 29 {9} GPs, 36 {11} CPs, and 33 {10} DMs. Each surviving individual was interviewed twice pre-exposure and twice post-exposure. All animals used were male and of sexual reproductive age, to comply with importation regulations. The experiments were run pre-exposure {post-exposure} on 3-26/5/2011 {3-9/8/2011}, 2-5/7/2012 {20-22/9/2012}, 6-11/7/2012 {8-10/11/2012} and 27-29/5/2013 {30/7/2013-2/8/2013} for GA, CP, DM and GP respectively.

### Data Analyses

We used four different methods to determine how the rodents perceive risk posed by the different snakes. First, we ran a Friedman’s test of concordance, comparing the way in which each rodent ranked the different snake species. The highest GUD received a rank of 1 and the lowest a rank of 3. We repeated the analysis on data for each individual pre-[post-] exposure.

To specifically address the low variation in GUDs in the pocket mice, we assessed activity patterns by running a log-linear tabulation analysis (multi way contingency table) on the proportion of foraged to unforaged trays. For each small mammal we compared the proportion of trays foraged based on snake treatment and experimental sequence (pre- and post-exposure)

Last, we averaged the GUD for each individual per snake treatment, resulting in one value for at each test sequence. We then ran in Systat13© a series of generalized linear models (GLM) using the mean GUD as the dependent variable. The first GLM used three independent variables; rodent species, (snake) treatments, and sequence. In addition, all two and three-way interactions between these variables were included. We did not use the full data set, but lowered the “noise” in the data by using the mean values. This normalization means that each individual provides two datapoints, one prior and one post exposure (too low for a meaningful comparison on the individual level). To increase the explanatory power for each species, we ran a GLM for each species individually as well. For the single species GLMs, we tested the independent variables: snake treatment and sequence (and the two-way interaction). Post-hoc pairwise comparisons were performed using Tukey’s Honestly Significant Differences (THSD) tests for variables that significantly affected variance. This analysis addresses a population-wise (or species-wise in this case) comparison for the broader differences and not in-population variation. We knowingly and purposefully removed individual ID for these reasons.

Last, we ran a random-forest Bayesian decision tree analysis in Statsoft Statistica©. This analysis best described as a categorical principal component analysis describes the importance of each variable, and category within each variable, in explaining the distribution of points of a dependent variable. Here we tested how the rodent’s GUD were distributed based on the species, the snake-treatment and the sequence of the measurement.

## RESULTS

At first encounter, all the species ranked the snakes similarly (Friedman’s test of concordance X_f_^2^ = 6.5, 2 df, p= 0.039, and *W* = 0.813), with lowest GUDs for the snakeless-control, higher GUDs for the evolutionarily novel snake, and still higher GUDs for the evolutionarily familiar snake. All rodent species perceived both snakes as threats (p= 0.003, 0.012 for known and sidewinders and horned vipers compared to control; Fig 1).

**Fig 1.**
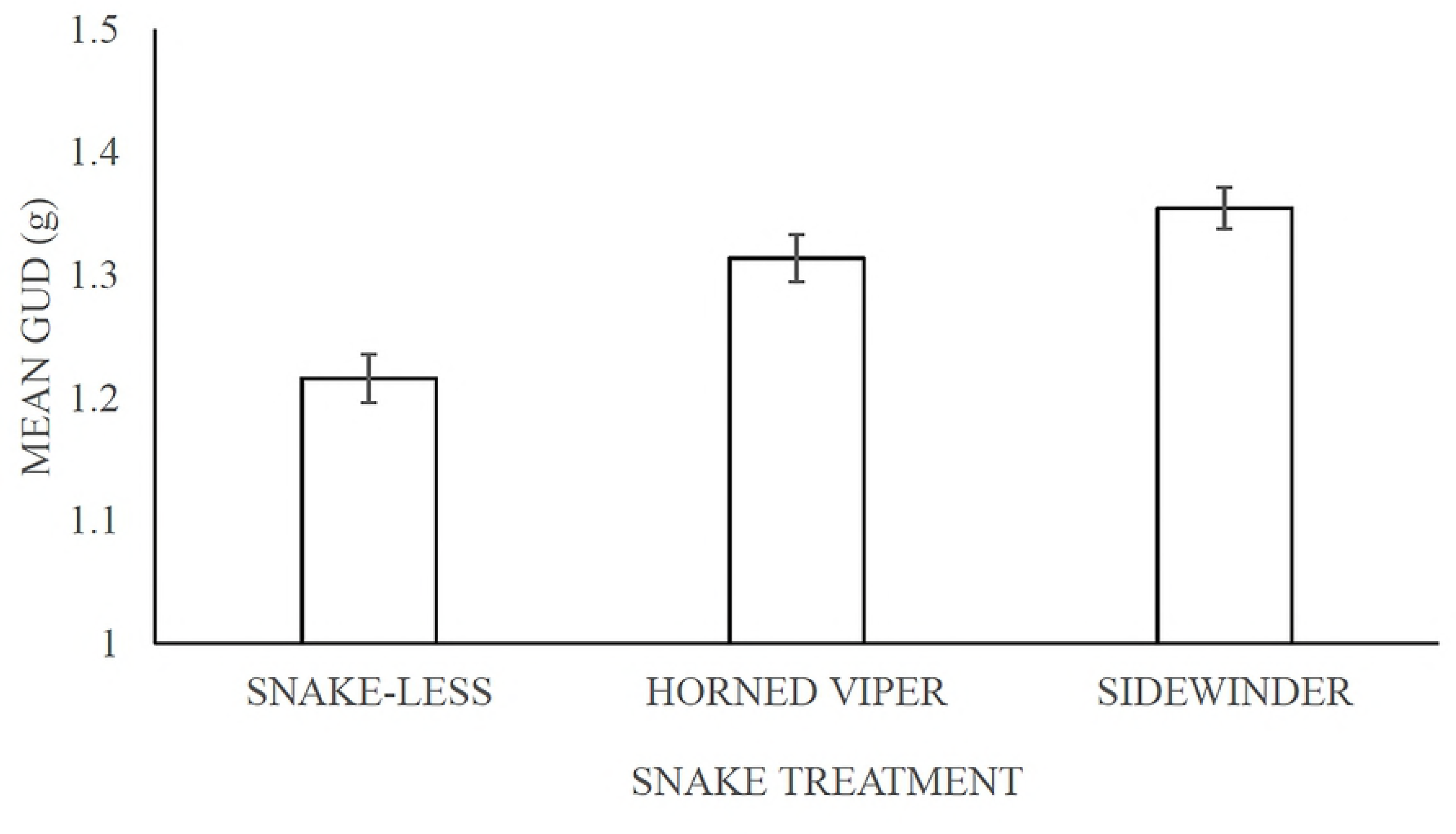
Giving-up densities (GUDs) ±SE combined for all four rodent species pre- and post-exposure.

Desert pocket mice showed increased GUDs in response to snake presence but did not distinguish between snake species in the magnitude of their GUDs. The remaining three rodents responded with 100% concordance, showing highest GUDs in response to their familiar snake, intermediate GUDs to the novel snake, and the lowest GUD when the cage contained no snake (*S2*). The non-significant difference between snakes in GP pre-exposure did not alter this finding. Post-exposure, the rodents showed complete concordance according to snake species: they all foraged least in the presence of the rattlesnake. (X_f_^2^ = 6.5, 2 df, p= 0.039, and *W*= 0.813).

**Table 1.**
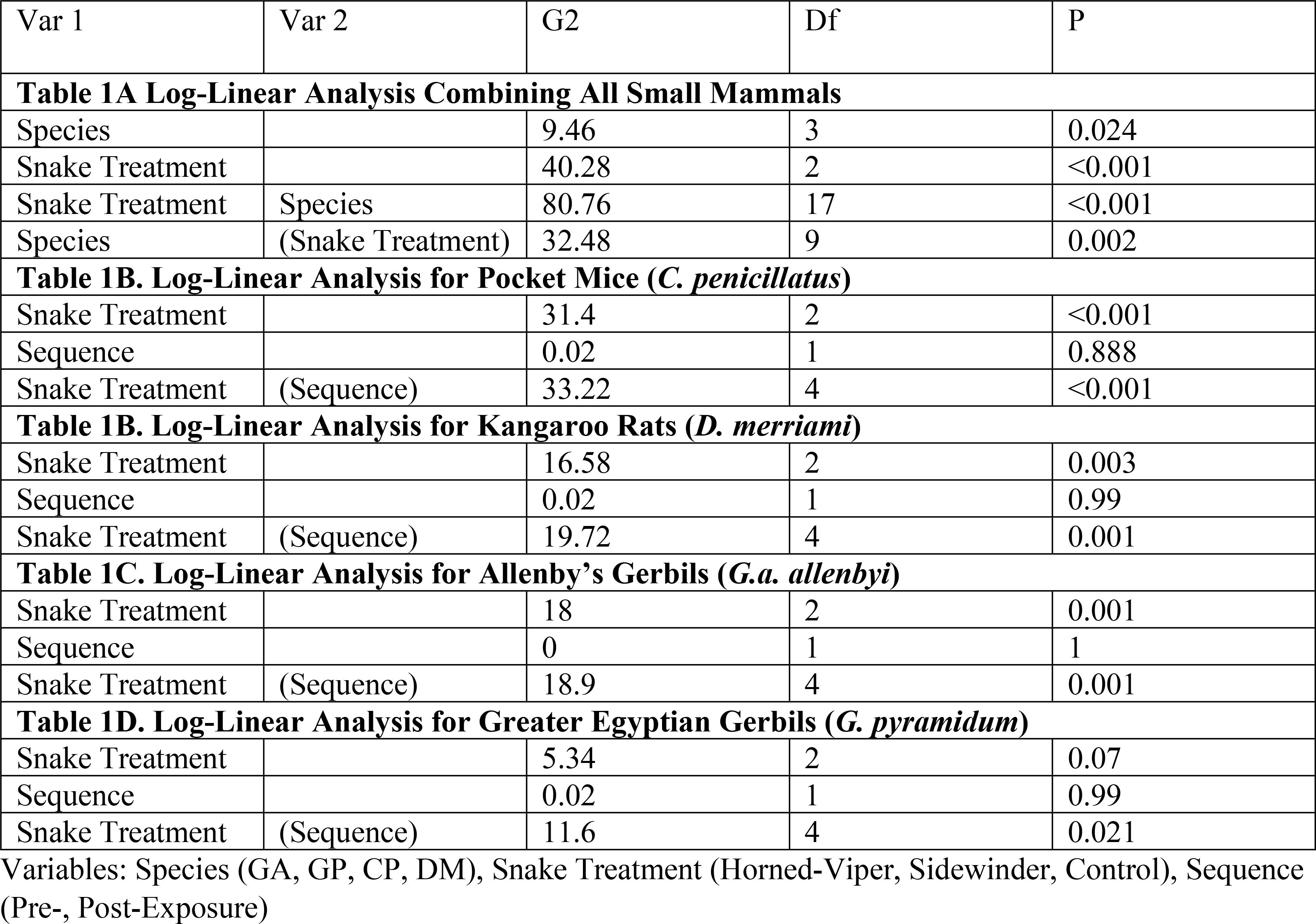
Log-Linear Analyses for all species combined (A) and each species separately (B-E)

Assessing the activity of each species of rodent, using proportion of patches of this treatment in which foraging activity took place, similar patterns emerge. We found that the species each exhibited different activity preferences (Table 1A). None of the rodents foraged a greater proportion of trays before the snake exposure than after. However, for all four rodents the exposure changed the willingness to forage in difference snake treatments (Tables 1 B-E). Pre-exposure, three species were active in more compartments with the novel snake than with the evolutionarily familiar one (Fig 2 A). Contrarily, GP investigated more compartments with the familiar snakes than novel snakes, and DM foraged in more compartments with novel snakes than in snake-less compartments. Post-exposure, all rodents foraged most in the snake-free-control over the snake treatments (Fig 2 B). Three of the four species foraged least in the compartment with sidewinder rattlesnakes. The GAs, foraged in more horned viper treatments than near sidewinders. GA’s activity pattern did not vary between pre- and post-exposure interviews.

**Fig 2.**
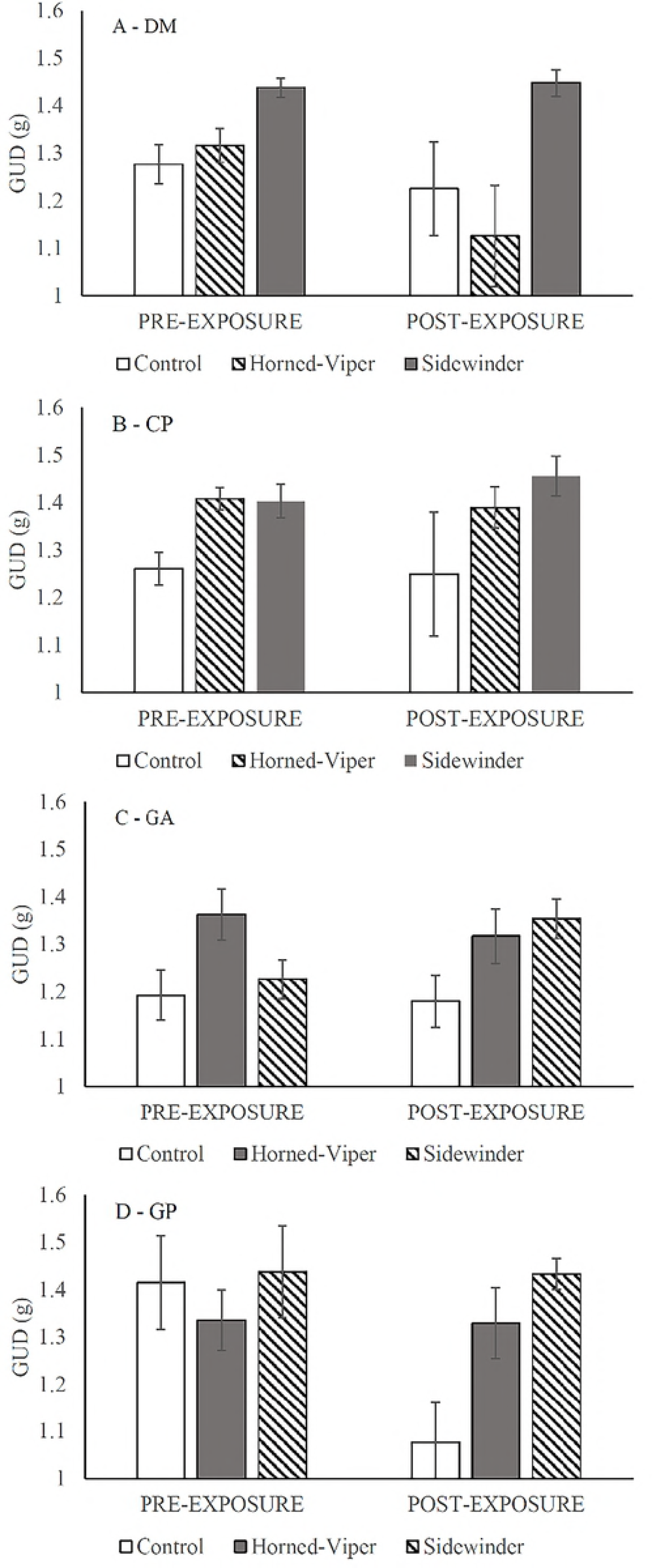
Cumulative proportion (active/total per treatment) of active foraging trays pre- (A) and post-exposure (B). “N” represents the novel snake treatment for each species. Black, gray and white bars represent sidewinder rattlesnakes, Saharan horned vipers and snake-less treatments respectively.

**Table 2.** ANOVA Tables for General Linear Models for all species combined (A) and each species separately (B-E)

| Var.1 | Var.2 | Var.3 | Type III SS | df | Mean Squares | F-Ratio | p-Value |
| --- | --- | --- | --- | --- | --- | --- | --- |
| **Table 2A. ANOVA Table for all four species N=582, Multiple R<sup>2</sup>=0.346** |  |  |  |  |  |  |  |
| Species |  |  | 0.591 | 3 | 0.197 | 2.741 | 0.043 |
| Sequence |  |  | 0.007 | 1 | 0.007 | 0.091 | 0.762 |
| Snake Treatment |  |  | 1.967 | 2 | 0.984 | 13.691 | <0.001 |
| Species | * Sequence |  | 0.165 | 3 | 0.055 | 0.766 | 0.513 |
| Snake Treatment | * Sequence |  | 0.793 | 2 | 0.397 | 5.52 | 0.004 |
| Species | *Snake Treatment |  | 0.973 | 6 | 0.162 | 2.256 | 0.037 |
| Snake Treatment | * Sequence | *Species | 0.305 | 6 | 0.051 | 0.708 | 0.643 |
| Error |  |  | 40.093 | 558 | 0.072 |  |  |
| **Table 2B. ANOVA table for Desert Pocket Mouse (*C. penicillatus*) N=141, Multiple R<sup>2</sup>=0.324** |  |  |  |  |  |  |  |
| Sequence |  |  | 0.001 | 1 | 0.001 | 0.03 | 0.863 |
| Snake Treatment |  |  | 0.584 | 2 | 0.292 | 6.16 | 0.003 |
| Snake Treatment | * Sequence |  | 0.026 | 2 | 0.013 | 0.272 | 0.762 |
| Error |  |  | 6.404 | 135 | 0.047 |  |  |
| **Table 2C. ANOVA table for Allenby's gerbil (*G. andersoni allenbyi*) N=210, Multiple R<sup>2</sup>=.225** |  |  |  |  |  |  |  |
| Sequence |  |  | 0.018 | 1 | 0.018 | 0.164 | 0.686 |
| Snake Treatment |  |  | 0.719 | 2 | 0.359 | 3.276 | 0.04 |
| Snake Treatment | * Sequence |  | 0.242 | 2 | 0.121 | 1.102 | 0.334 |
| Error |  |  | 22.383 | 204 | 0.11 |  |  |
| **Table 2D. ANOVA table for Greater Egyptian Gerbil (*G. pyramidum*) N=104, Multiple R<sup>2</sup>=.460** |  |  |  |  |  |  |  |
| Sequence |  |  | 0.008 | 1 | 0.008 | 0.146 | 0.703 |
| Snake Treatment |  |  | 0.79 | 2 | 0.395 | 6.926 | 0.002 |
| Snake Treatment | * Sequence |  | 0.695 | 2 | 0.347 | 6.091 | 0.003 |
| Error |  |  | 5.476 | 96 | 0.057 |  |  |
| **Table 2E. ANOVA table for Merriam's Kangaroo Rat (*D merriami*) N=129, Multiple R<sup>2</sup>=.406** |  |  |  |  |  |  |  |
| Sequence |  |  | 0.138 | 1 | 0.138 | 2.912 | 0.09 |
| Snake Treatment |  |  | 0.891 | 2 | 0.446 | 9.4 | <0.001 |
| Snake Treatment | * Sequence |  | 0.162 | 2 | 0.081 | 1.709 | 0.185 |
| Error |  |  | 5.83 | 123 | 0.047 |  |  |

Variables: Species (GA, GP, CP, DM), Snake Treatment (Horned-Viper, Sidewinder, Control), Sequence (Pre-, Post-Exposure)

The GLM combining all four species showed that each species foraged differently in the interview chambers (Table 2). The heteromyids foraged less than the gerbils. The pocket mice (CP), and kangaroo rats (DM) foraged to a mean GUDs (±SE) of 1.34±0.019g and 1.32±0.019g, respectively. Allenby’s gerbils (GA) and the Egyptian gerbils (GP) foraged to mean GUD of 1.24±0.02g and 1.29±0.027g, respectively. In response to the snake treatments, the rodents overall foraged least in the presence of the rattlesnake, and most in the control treatment (Fig 3, Sup. Appendix 2). Post-hoc pairwise comparison (THSD) found a significant difference between, the control and horned viper (p=0.009), the control and sidewinder (p<0.001), and control and between the horned viper and sidewinder (p=0.006). After two months of exposure, all four species exhibited a similar trend of decreased foraging in the presence of the sidewinder rattlesnake (Fig 3) as shown in each of the single species models (Table 2).

**Fig 3.**
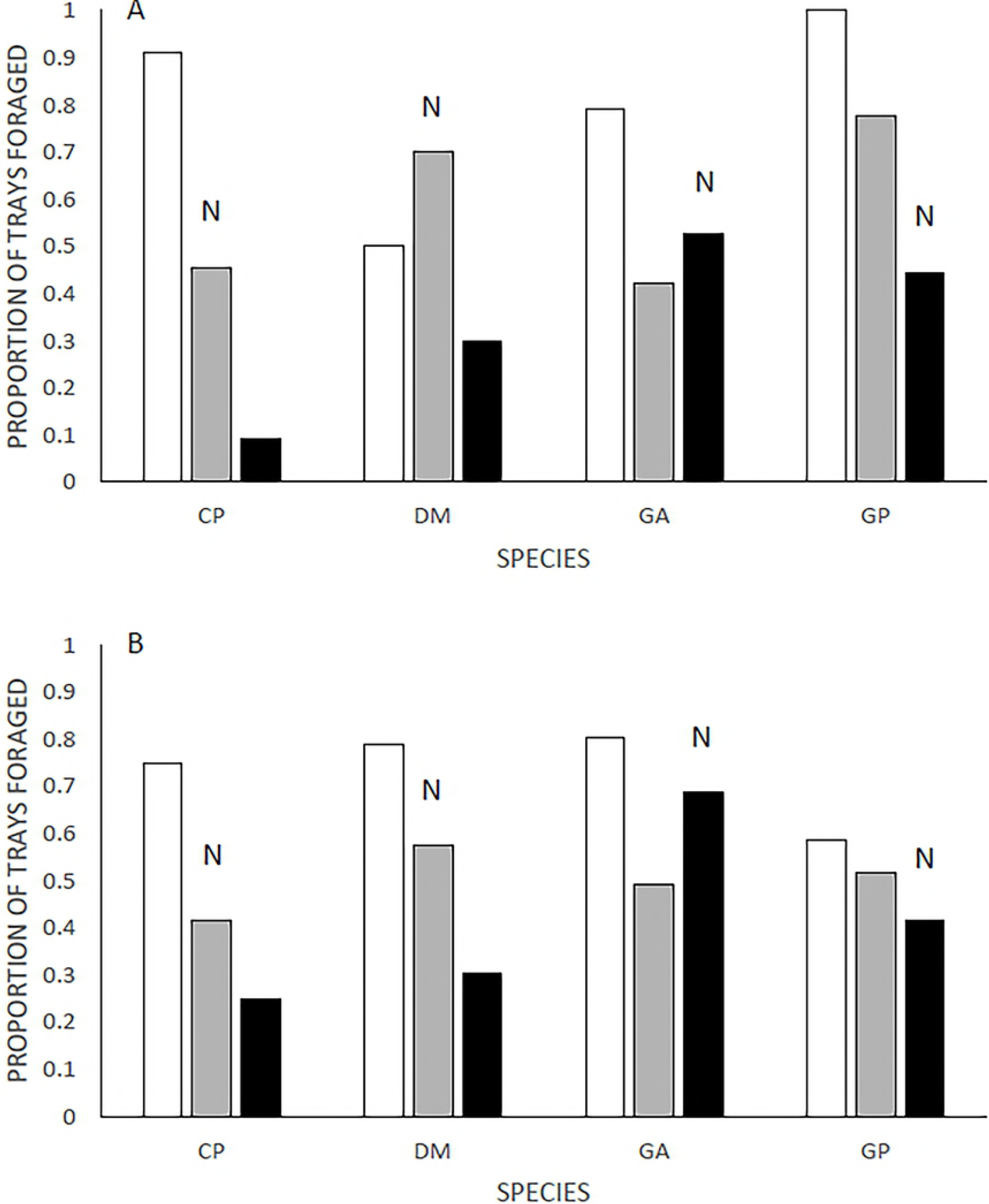
Perception of risk of familiar and novel snakes, in pre- and post-exposure interviews, as reflected by giving-up densities (GUDs) ±SE. Each frame subfigure depicts the response of one species: (A) DM, (B) CP, (C) GA and (D) GP. The rodents were exposed to Saharan horned vipers from the Negev and sidewinder rattlesnakes from the Mojave. In each frame the diagonal striped bar reflects the snake that is evolutionarily novel, the gray bar is the known snake and the white bar is the snake-less control.

The random forest analysis resulted in a model with mid-range risk-estimates (±SE) of 0.09±0.02 and 0.11±0.03 for the training and model respectively. The model confirmed strong species and snake treatment effects (importance of 0.993 and 1.0 out of 1.0 respectively) and suggested lower importance of 0.326 for the sequence. These values explain the rate of decisions each variable affected. This analysis unexpectedly separated GA from the other species at the first split (Fig 4; Supplement 3). The higher a split (further left) the stronger greater the accuracy of that split, thus more credible. For the GAs the difference between the control and snakes was critical, and the model suggests that the novel pit viper was assessed as slightly riskier post exposure. The GUDs reflect the change in perception with means of 1.27±0.06 g and 1.29±0.06 g pre- and post-exposure respectively.

**Fig 4.**
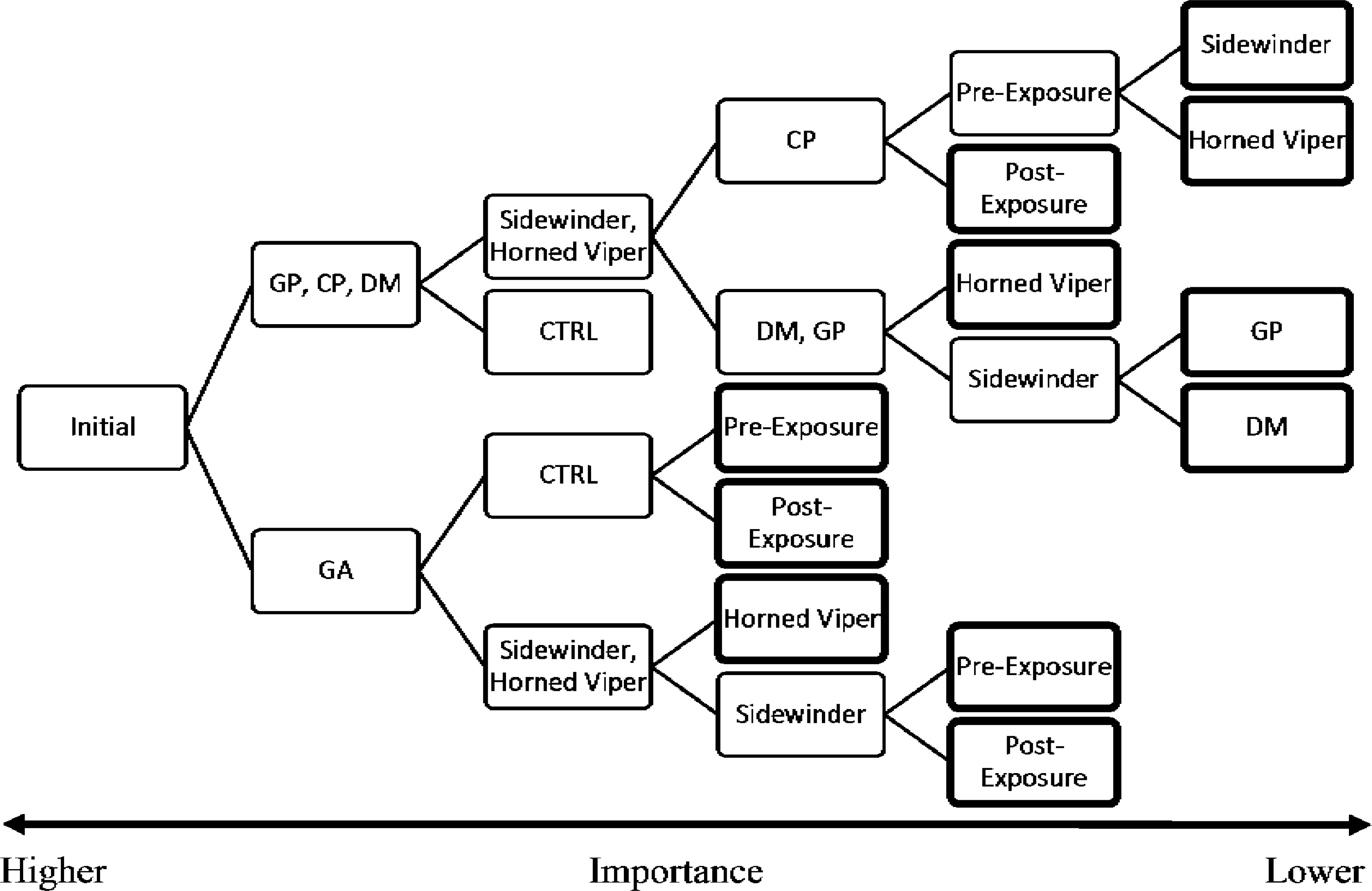
Random-Forest Decision-Tree with GUD as the dependent variable and species, snake-treatment and sequence as the independent variables. The figure is read from left to right with greater value to the initial nodes (left) than to final nodes marked with a bold outline.

The difference in variance between the GUDs in the control for all three other species (CP, DM & GP) was too small for the model to predict divergence between them. However, in the response to the snakes a clear divergence between CP and the larger rodents was found. The Pocket mice initially foraged less near the novel snake, but post exposure avoided both snakes equally. The model does not predict change in the larger rodents assessment of risk from the snakes after exposure, but clumps these points together to set aside the sidewinder as greater risk for both. The mean GUD for both species combined in the presence of the horned vipers was 1.3±0.05g. The difference between the response to each snake type was larger in the kangaroo rats (DM), with a GUD of 1.45±0.03g in the presence of the sidewinders, than in the gerbils (GP), with respective GUDs of 1.34±0.08 g.

## DISCUSSION

All rodents began by assessing the snake with which they share evolutionary history as an equal or greater threat to the novel snake. However after two months of interacting with uncaged sidewinders and horned vipers, all four rodents ranked the heat-sensing sidewinder as the greater threat. We chose to structure this discussion according to three major comparisons: two intracontinental comparisons (within families) and one intercontinental comparison between the gerbils and heteromyids.

### Heteromyids

Pocket mice and kangaroo rats provide examples of opposite strategies in managing risk from snakes (as shown here), and other predators such as owls [28,32]. Why the stark difference? We do not believe it has to do solely with size, but in variation in anti-predator adaptations. The first reason behind this speculation is a number of studies investigating microhabitat selection in the kangaroo rats and pocket mice of different species [33–35]. In those studies, the kangaroo rats would use both bush and open microhabitat whilst pocket mice were particular to the bush. The interpretation these studies gave are based in both locomotion and signaling resulting from the kangaroo rats’ bipedal agility. On the opposite side the evolutionary strength of the pocket mice is attributed to torpor which they apply to minimize risk and survive harsh weather events. In addition, the kangaroo rats are able to ward off snakes using warning signals, foot drumming and kicking sand in the face of their predators [25,36]. In facing a striking snake, they are capable of hopping backwards [37–39] and to heights exceeding 2 m [40]. In contrast, the pocket mice remain bush-bound, and avoid predators by climbing into dense vegetation and are presumed to apply a torpor mechanism to reduce dependency on the foraging when risk levels are too high [33,35].

In the interview chambers, differences among the species were well represented. The pocket mice avoided risk where possible, leaving high GUDs near snakes, affected by the 70-80% of rooms unforaged. The avoidance of the snakes was strongly offset by significantly lower GUD and high activity in the snake-free control rooms. Meanwhile, the kangaroo rats exhibited an inquisitive nature seen by the high proportion of foraged rooms. The genus *Dipodomys* (kangaroo rats) has morphological adaptation that allow them to both located predators and avoid their attack. Inflated auditory bullae allow the kangaroo rats to hear predators, such as owls, approaching from a large distance [34,41,42]. The kangaroo rat’s powerful hind-legs allow them to hop out of harm’s way to heights above one meter and are able to change the direction of movement using tail flicks while in the air [37]. Thanks to their morphological adaptations, kangaroo rats are able of greater risk-taking than the pocket mice [43]. This risk-taking behavior, verging on being dare-devils, is best exhibited by the increased resource use and patch activity in the treatment with the novel snake (greater than the control). These strong differences in anti-predator adaptations, both behavioral and physical, are likely the evolutionary mechanism that allows for these species to coexist in the great basin deserts.

### Gerbils

The competition between GA and GP is a major model system for the study of the roles of competition, predation risk and parasitism in community structure. It the behavioral differences between these gerbils that allow them to coexist. They differ in habitat preference [44–46], in the time of night they are active [47,48], in the way they respond to different types of predators (snakes, owls) [49–52] and in the way they respond to inter- and intra-specific competition [53–55]. Surprisingly, despite those well documented behavioral differences, we found the species responded to the snakes in remarkably similar patterns.

Why did we find such similar patterns? The most likely explanation is that our systems were devoid of environmental heterogeneity. During the exposure period, we found species-specific—spatially explicit—responses to the distribution of risk posed by each snake and in combination with barn owls [28,30,32,56]. However, in the enclosed systems, where individual gerbils forage without competition, the response of both species to the risk of predations is similar.

This experiment revealed that the gerbils were attentive to the type of predation-risk present and their response to that risk is relatively plastic. Pre-exposure, both gerbils recognized the novel sidewinder, as a risk (higher GUDs than the control) but not as great a risk as the known horned viper. The change in perceived risk towards the novel sidewinder suggests the gerbils gained information about the new predator. Post-exposure, the mean GUDs being similar for both snake species, suggests the gerbils were able to learn, in the minimum, that these new predators are snakes. Despite both having some changes in their response the GAs exhibit a stronger tendency to adapt to the novel risk (based on the random-forest), not surprising for a species that is known for balancing the risk from predators with stronger competitors, *a.k.a* a crumb picking foraging strategy. The GA’s are known to assess GP (and *Gerbillus gerbillus*) activity and shift their foraging patterns to exploit patches more thoroughly when these dominant species are around [47,48].

Another possible explanation is delayed response to stimulus. In neurological studies delayed response to a novel threat is commonly studied in contexts of neophobia and clasical conditioning. In these types of studies lab mice, rats and rabbits are taught to recognize a novel object, sound, or image as a predation-cue [57,58]. Intrinsically, most rodents fear novel objects, but do not innately respond to them to the extent of the danger that they “actually” pose. In many cases, they remain naive to the proper response to these novel threats [59,60]. Despite being naïve to the dangers of the sidewinder rattlesnake at the start of the experiments, both gerbils quickly learned to respond to the snakes and both rank them as a risk. In out measurements in the aviary they both ranked snakes as a lower risk than (lower GUDs) than owls [28,30,32]. The results of the comparison between the gerbils highlight the importance of competition to species that have less spatial segregation than the North American heteromyids [44,47].

### Intercontinental Wide Consequences

During the pre-exposure interviews all rodents feared their evolutionarily familiar snake equally or more than the novel one. In particular, gerbils showed higher GUDs in response to greater Saharan horned vipers, and the heteromyids showed higher GUDs in response to the sidewinder rattlesnake. This coincides with the snake species that each species evolved with. However, this may also reflect the predator to which each of the rodents has individually been exposed to previously since all animals in these experiments were wild-caught. Overall, the gerbelline species were willing to take more risk investigating the predators, while the heteromyids preferred to avoid both species. This reluctance to take risk in the heteromyids, suggests an overall greater “respect” to risk posed by snakes, possibly due to their evolution alongside snakes that have heat-sensing capabilities [32].

All four rodents, showed an ability to varying extent, to differentiate between the snake species and to categorize the heat-sensing sidewinder as a greater threat post-exposure. This could be attributed to two explanations. First, the rodents may have learned to identify the musk produced by each species, as kangaroo rats are known to do [36]. Second, given the dark conditions, aimed to highlight the difference between the snakes, the rodents may have been responding most strongly to the sidewinders as they were more active in their cages.

The fur-lined cheek pouches, hypothesized to give the heteromyids an advantage in maximizing harvest, under similar risk conditions, did not appear to function in such a manner in the presence of the snakes [32]. The greater sensitivity of heteromyids to the interaction with the snakes meant that despite having the ability to forage quickly in the presence of the predators, the heteromyids simply avoided these patches resulting in higher giving up densities than their gerbilline counterparts.

Perhaps the most interesting aspect of this comparison was the broader examination of the convergence of the two deserts’ rodent and predator communities. Despite the physical and dietary similarities, we found a large number of key differences in the way our four species strategized in response to snake predators. Our experiment shows a tendency of North American species to focus on predation risk while the gerbil responses to the caged snakes were more plastic, likely suggesting other elements come to play in addition to snakes in that system. Why are the heteromyids more sensitive to the risk posed by snakes? The only likely explanation is the one that made us choose these systems for comparison, *i.e*. the evolution along-side predators with a lethal weapon—the heat sensing pits. While sidewinders were the clear choice in terms of physical convergence with the horned vipers, they are only one of 13 species of rattlesnakes that call the Great Basin deserts home, and all possess the infra-red sensory ability [61]. In comparison, there are only five vipers in the Negev and the Sahara, and they all are blind on moonless nights [62]. The high diversity of lethal predators in North America suggests the pressure to manage the risk from snakes has been a lot more important in the evolution of heteromyids. From that importance stems their sensitivity and acuity to the presence and activity patterns of the snakes they encounter.

To conclude we can recapitulate the answers we found to each of our study questions. (1) Middle Eastern gerbils responded less to predation risk posed by snakes than their North American convergent counterparts, kangaroo rats and pocket mice. (2) At first encounter, both kangaroo rats and gerbils recognized the novel snake as a lesser (or equal) risk to that of their familiar snakes. Pocket mice avoided both snakes equally. (3) Post-exposure, gerbils and pocket mice assessed both snake species as similarly dangerous. However, kangaroo rats rank the novel, horned viper as a lesser threat than the known heat-sensing sidewinder.

## ACKNOWLEDGMENTS

We would like to acknowledge the US-Israel Binational Science Foundation (BSF) for funding this project (BSF-2008163). The permits for this project were obtained from Ben Gurion University of the Negev Ethics in Animal Research Committee (Permit IL-73-11-2009). The permits for animal shipping, handling and experimentation were obtained from the Israel Nature and National Parks Authority (INPA) (permits 2011/38131 and 2012/12524). We would also like to acknowledge the assistance in obtaining the research populations, field work and training of A. Bouskila, P. C. Rosen, D. Burns, J. R. St. Juliana, C. Downs, E. Kiekebusch, S. Summerfield and I. Hoffmann. We thank. C. Hammond for the illustrations. This is publication number XXX of the Mitrani Department for Desert Ecology.

## LITERATURE CITED

1. Chew RM, Eastlake Chew A. Energy Relationships of the Mammals of a Desert Shrub (*Larrea tridentata*) Community. Ecol Monogr. 1970;40: 1–21.

2. Rosenzweig ML, Winakur J. Population ecology of desert rodent communities : habitats and environmental complexity. Ecology. 1969;50: 558–572.

3. Brown JH, Kelt DA, Fox BJ, The S, Naturalist A, December N. Assembly Rules and Competition in Desert Rodents. Am Nat. 2002;160: 815–818.

4. Kotler BP. Risk of predation and the structure of desert rodent communities. Ecology. 1984;65: 689–701.

5. Brown JS. Desert rodent community structure: A test of four mechanisms of coexistance. Ecol Monogr. 1989;59: 1–20.

6. Meserve PL, Dickman CR, Kelt D a. Small mammal community structure and dynamics in aridlands: overall patterns and contrasts with Southern Hemispheric systems. J Mammal. 2011;92: 1155–1157. doi:10.1644/11-MAMM-S-186.1

7. Mares MA. Desert Rodents, Seed Consumption, and Convergence. Bioscience. 1993;43: 372–379. doi:10.2307/1312045

8. Houseman GR, Mittelbach GG, Reynolds HL, Gross L. Perturbations Alter Community Convergence, Divergence, and Formation of Multiple Community States. Ecology. 2008;89: 2172–2180.

9. Trontelj P, Blejec A, Fiser C. Ecomorphological convergence of cave communities. Evolution (N Y). 2012;66: 3852–3865. doi:10.1111/j.

10. Wassenaar TD, van Aarde RJ, Pimm SL, Ferreira SM. Community Convergence in Disturbed Subtropical Dune Forests. Ecology. 2005;86: 655–666.

11. Davidson DW, Inouye RS, Brown JH. Granivory in a desert ecosystem: experimental evidence for indirect facilitation of ants by rodents. Ecology. 1982;65: 1780–1786.

12. Brown JH, Munger JC. Experimental manipulation of a desert rodent community : food addition and species removal. Ecology. 1985;66: 1545–1563.

13. Brown JS, Kotler BP, Mitchell WA. Foraging theory, patch use, and the structure of a Negev granivore community. Ecology. 1994;75: 2286–2300.

14. Ori C. Crotalus cerastes. In: Animal Diversity Web [Internet]. 2000. Available: http://animaldiversity.ummz.umich.edu/accounts/Crotalus_cerastes/

15. Anderson I. Cerastes cerastes. In: Animal Diversity Web. [Internet]. 2011. Available: http://animaldiversity.ummz.umich.edu/accounts/Cerastes_cerastes/

16. Brown JS, Kotler BP, Valone TJ. Foraging under Predation: a Comparison of Energetic and Predation Costs in Rodent Communities of the Negev and Sonoran Deserts. Aust J Zool. 1994;42: 435–448.

17. Kotler BP, Brown JS. Environmental heterogeneity and the coexistance fo desert rodents. Annu Rev Ecol Evol Syst. 1988;19: 281–307.

18. Rosenzweig ML, McCord RD. Incumbent replacement: evidence for long-term evolutionary progress. Paleobiology. 1991;17: 202–213.

19. Vincent TL, Brown JS. Evolutionary game theory, natural selection, and Darwinian dynamics. Cambridge, UK: Cambridge University Press; 2005.

20. Wüster W, Peppin L, Pook CE, Walker DE. A nesting of vipers: Phylogeny and historical biogeography of the Viperidae (Squamata: Serpentes). Mol Phylogenet Evol. Elsevier Inc.; 2008;49: 445–59. doi:10.1016/j.ympev.2008.08.019

21. Pappas TC, Motamedi M, Christensen BN. Unique temperature-activated neurons from pit viper thermosensors. Am J Physiol Cell Physiol. 2004;287: C1219–28. doi:10.1152/ajpcell.00040.2004

22. Goodfriend W, Ward D, Subach A. Standard operative temperatures of two desert rodents, *Gerbillus allenbyi* and *Gerbillus pyramidum*: The effects of morphology, microhabitat and environmental factors. J Therm Biol. 1991;16: 157–166. doi:10.1016/0306-4565(91)90038-4

23. Lancaster E. Dipodomys merriami. In: Animal Diversity Web [Internet]. 2000. Available: http://animaldiversity.ummz.umich.edu/accounts/Dipodomys_merriami/

24. Chebes L. Cheatodipus penicillatus. In: Animal Diversity Web [Internet]. 2002 [cited 1 Jan 2017]. Available: http://animaldiversity.org/accounts/Chaetodipus_penicillatus/

25. Randall J. Species-specific footdrumming in kangaroo rats: *Dipodomys ingens, D. deserti, D. spectabilis*. Anim Behav. 1997;54: 1167–75. Available: http://www.ncbi.nlm.nih.gov/pubmed/9398370

26. Webber MM, Jezkova T, Rodríguez-Robles JA. Feeding Ecology of Sidewinder Rattlesnakes, *Crotalus cerastes* (Viperidae). Herpetologica. 2016;72: Herpetologica-D-15-00031. doi:10.1655/Herpetologica-D-15-00031

27. Bleicher SS. Heat and Humidity Alter Predation Cues in Gerbillus andersoni allebyi. MS Thesis. Ben Gurion University of the Negev. Ben Gurion University of the Negev. 2012.

28. Bleicher SS. Divergent behaviour amid convergent evolution: common garden experiments with desert rodents and vipers. Ph.D. Disertation, Univeristy of Illinois. University of Illinois at Chicago. 2014.

29. Bleicher SS, Dickman CR. Bust economics: foragers choose high quality habitats in lean times. PeerJ. 2016;1: 1–15. doi:10.7717/peerj.1609

30. Bleicher SS, Brown JS, Embar K, Kotler BP. Novel predator recognition by Allenby’s gerbil (*Gerbillus andersoni allenbyi*): do gerbils learn to respond to a snake that can “see” in the dark? Isr J Ecol Evol. 2016;62: 178–185. doi:10.1080/15659801.2016.1176614

31. Brown JS. Patch use as an indicator of habitat preference, predation risk, and competition. Behav Ecol Sociobiol. 1988;22: 37–47. doi:10.1007/BF00395696

32. Kotler BP, Brown JS, Bleicher SS, Embar K. Intercontinental-wide consequences of compromise-breaking adaptations: the case of desert rodents. Isr J Ecol Evol. 2016;62: 186–195.

33. Thompson SD. Microhabitat Utilization and Foraging Behavior of Bipedal and Quadrupedal Hetermoyid Rodents. Ecology. 1982;63: 1303–1312.

34. Kotler BP, Brown JS, Smith RJ, Wirtz WOI. The effects of morphology and body size on rates of owl predation on desert rodents. Oikos. 1988;53: 145–152.

35. Lemen CA, Rosenzweig ML. Microhabitat Selection in Two Species of Heteromyid Rodents. Oecologia. 1978;33: 127–135.

36. Bouskila A. Interactions between predation risk and competition : a field study of kangaroo rats and snakes. Ecology. 1995;76: 165–178.

37. Bartholomew GA, Caswell HHJ. Locomotion in Kangaroo Rats and Its Adaptive Significance. J Mamm Evol. 1951;32: 155–169.

38. Randall JA, Stevens CM. Footdrumming and Other Anti-Predator Responses in the Bannertail Kangaroo Rat (*Dipodomys spectabilis*). Behav Ecol Sociobiol. 1987;20: 187–194.

39. Clark RW, Dorr SW, Whitford MD, Freymiller GA, Hein SR. Comparison of antisnake displays in the sympatric desert rodents *Xerospermophilus tereticaudus* (round-tailed ground squirrels) and *Dipodomys deserti* (desert kangaroo rats). J Mammal. 2016;97: 1–9. doi:10.1093/jmammal/gyw137

40. Merlin P. Heteromyidae: Kangaroo Rats & Pocket Mice. In: Arizona-Sonora Desert Museum [Internet]. 2014 [cited 6 Jan 2017]. Available: https://www.desertmuseum.org/books/nhsd_heteromyidae.php

41. Webster DB, Webster M. The specialized auditory system of kangaroo rats. Contrib Sens Physiol. 1984;8: 161–196.

42. Longland WS, Price M V. Direct observations of owls and heteromyid rodents : Can predation risk explain microhabitat use ? Ecology. 1991;72: 2261–2273.

43. Rosenzweig ML. Habitat selection experiments with a pair of coexisting Heteromyid rodent species. Ecology. 1973;54: 111–117.

44. Kotler BP, Brown JS, Oldfield A, Thorson J, Cohen D. Foraging substrate and escape substrate: patch use by three species of gerbils. Ecology. 2001;82: 1781–1790. doi:10.1890/0012-9658(2001)082[1781:FSAESP]2.0.CO;2

45. Abramsky Z, Rosenzweig ML, Pinshow B. The shape of a gerbil iosocline measured using principles of optimal habitat selection. Ecology. 1991;72: 329–340.

46. Kotler BP, Brown JS, Mitchell WA. Environmental factors affecting patch use in two species of gerbelline rodents. J Mammal. 1993;74: 614–620. doi:10.2307/1382281

47. Kotler BP, Brown JS, Subach A. temporal foragers : of optimal coexistence of species mechanisms of sand dune gerbils by two species partitioning. Oikos. 1993; 548–556.

48. Kotler BP, Brown JS, Dall SRX, Gresser S, Ganey D, Bouskila A. Foraging games between gerbils and their predators : temporal dynamics of resource depletion and apprehension in gerbils. Evol Ecol Res. 2002;4: 495–518.

49. Embar K, Kotler BP, Mukherjee S. Risk management in optimal foragers: the effect of sightlines and predator type on patch use, time allocation, and vigilance in gerbils. Oikos. 2011;120: 1657–1666. doi:10.1111/j.1600-0706.2011.19278.x

50. Abramsky Z, Strauss E, Subach A, Riechman A, Kotler BP. The effect of barn owls (*Tyto alba*) on the activity and microhabitat selection of *Gerbillus allenbyi and G. pyramidum*. Oecologia. 1996;105: 313–319. doi:10.1007/BF00328733

51. Kotler BP, Brown JS, Slotow RH, Goodfriend WL, Strauss M. The influence of snakes on the foraging behavior of gerbils. Oikos. 1993;67: 309–316.

52. Kotler BP, Blaustein L, Dednam H. The specter of predation: the effects of vipers on the foraging behavior of two Gerbilline rodents. Isr J Zool. 1993;39: 11–21. doi:10.1080/00212210.1993.10688690

53. Berger-Tal O, Embar K, Kotler BP, Saltz D. Everybody loses : intraspecific competition induces tragedy of the commons in Allenby’s gerbils. Ecology. 2015;96: 54–61.

54. Ovadia O, Abramsky Z, Kotler BP, Pinshow B. Inter-specific competitors reduce inter-gender competition in Negev Desert gerbils. Oecologia. 2005;142: 480–8. doi:10.1007/s00442-004-1726-9

55. Abramsky Z, Rosenzweig ML, Subach A. The cost of interspecific competition in two gerbil species. J Anim Ecol. 2001;70: 561–567.

56. Embar K, Raveh A, Hoffmann I, Kotler BP. Predator facilitation or interference: a game of vipers and owls. Oecologia. 2014;174: 1301–9. doi:10.1007/s00442-013-2760-2

57. Bevins RA, Besheer J, Palmatier MI, Jensen HC, Pickett KS, Eurek S. Novel-object place conditioning: Behavioral and dopaminergic processes in expression of novelty reward. Behav Brain Res. 2002;129: 41–50. doi:10.1016/S0166-4328(01)00326-6

58. Weiss C, Disterhoft JF. Eyeblink conditioning and novel object recognition in the rabbit: Behavioral paradigms for assaying psychiatric diseases. Front Psychiatry. 2015;6: 1–9. doi:10.3389/fpsyt.2015.00142

59. Carthey AJR, Banks PB. Naïveté in novel ecological interactions: lessons from theory and experimental evidence. Biol Rev Camb Philos Soc. 2014;89: 932–49. doi:10.1111/brv.12087

60. Banks PB, Dickman CR. Alien predation and the effects on multiple levels of prey naivite. Trends Ecol Evol. 2007;22: 229–230. doi:10.1016/j.tree.2007.02.003

61. Fowlie JA. The snakes of Arizona, their derivation, distribution, description, and habits; a study in evolutionary herpetozoogeographic phylogenetic ecology. Fallbrook, CA: Azul Quinta Press; 1965.

62. Joger U. The venomous snakes of the Near and Middle East. Wiesbaden: L. Reichert; 1984.

